# *De novo* Design of a Polycarbonate Hydrolase

**DOI:** 10.1101/2023.03.10.532063

**Authors:** Laura H. Holst, Niklas G. Madsen, Freja T. Toftgård, Freja Rønne, Ioana-Malina Moise, Evamaria I. Petersen, Peter Fojan

**Affiliations:** Material Science and Engineering Group, Department of Materials and Production, Aalborg University, Denmark

## Abstract

Enzymatic degradation of plastics is currently limited to the use of engineered natural enzymes. As of yet, all engineering approaches applied to plastic degrading enzymes retain the natural *α*/*β* -fold. While mutations can be used to increase thermostability, an inherent maximum likely exists for the *α*/*β* -fold. It is thus of interest to introduce catalytic activity toward plastics in a different protein fold to escape the sequence space of plastic degrading enzymes. Here, a method for designing highly thermostable enzymes that can degrade plastics is described. This has been used to design an enzyme that can catalyze the hydrolysis of polycarbonate, which no known natural enzymes can degrade. Rosetta enzyme design is used to introduce a catalytic triad into a set of thermostable scaffolds. Through computational evaluation, a potential PCase was selected and produced recombinantly in *E. coli*. CD spectroscopy suggests that the design has a melting temperature of >95°C. Activity towards a commercially used polycarbonate (Makrolon 2808) was confirmed using AFM, which showed that a PCase had been designed successfully.

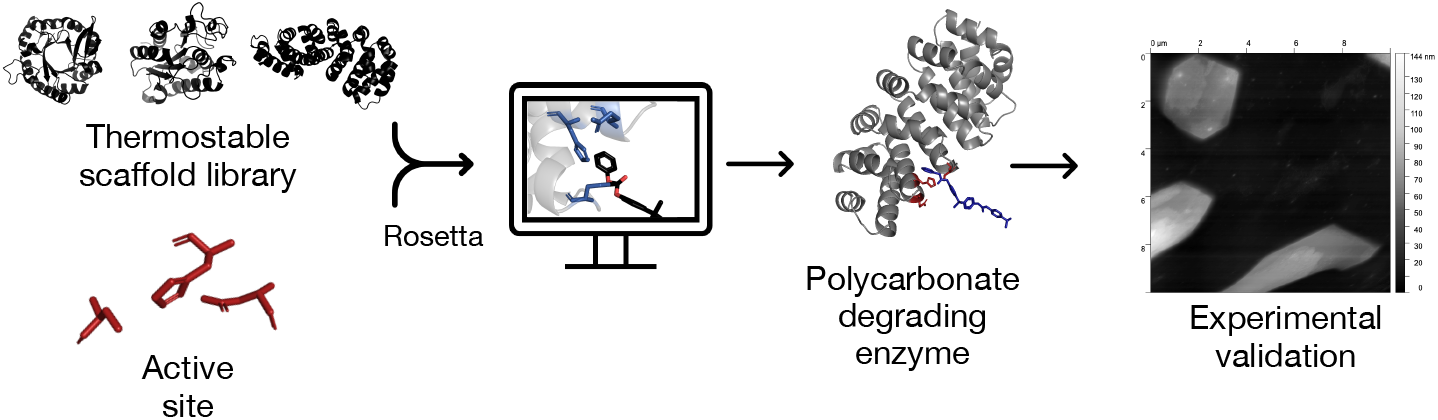

## INTRODUCTION

More than 350 million tons of plastic is produced each year, and only around 10% of plastic waste is recycled [1, 2]. The main recycling process used today is based on the mechanical break down of the polymers, which causes a decline in the quality of the material. If instead plastic is depolymerized to monomers, which could be reused as building blocks in synthesis of new plastics, the material properties would be maintained. [1] This goal can be reached by chemical degradation, however, it is an unfavorable process in terms of energy consumption since it typically happens at high temperatures and pressure. Enzymatic degradation could on the other hand be an advantageous alternative since the depolymerization of the plastics would occur under mild conditions. Since fossil fuel based polymers are not of natural origin, it is difficult to find enzymes in nature that can break down plastics. [3] Until now only a few enzymes have been identified to degrade polyethylene terephthalate (PET) and polyurethane (PUR) or show activity towards polyamide (PA) and polyethylene (PE) [4, 5, 6]. The major approaches that have been used to identify new plastic depolymerizing enzymes typically include enrichment cultivation [7], metagenomics [8], and mining of genome databases [9, 10]. Since the identification of T*f*H cutinase from *Thermobifida fusca* [11], the first hydrolase to be reported active against PET, the main target in the field of enzymatic plastic degradation has been PET [12]. However, there is a need for enzymes, which can degrade plastics apart from PET since it only constitutes *∼*6% of the global plastic production. [2, 12]

The major challenge to be addressed by enzymatic degradation of plastic is the high level of crystallinity. Within crystalline patches, the polymer chains exhibit low chain flexibility, which hinders the enzyme from attaching to the individual polymer chains. [13] In order to generate optimal interaction between the enzyme and the polymer, the process conditions and the enzyme need to be optimized. It is known that chain flexibility of the polymer increases as the temperature approaches the glass transition temperature. Chain flexibility is also increased by the presence of organic solvents. As a result, the degradation efficiency may increase if the reaction is performed at higher temperature or with the addition of organic solvents, requiring higher stability of the enzymes. [14, 13, 12]

All enzymes identified to degrade PET belong to the *α*/*β*-fold family [15]. The enzymes are typically classified as cutinases, lipases, or esterases [6]. In 2009, a cutinase from *Humicola insolens* (HiC) was used to degrade 97% of a low crystallinity PET film within 96 hours when incubated at 70°C [16]. In 2012, a cutinase was isolated from leaf-branch compost, LCC, using a metagenomics approach [8]. LCC could degrade *∼*25% of a commercial PET sample in 24 hours at 70°C [17]. In 2016, the mesophilic *Is*PETase enzyme was identified in *Ideonella sakaiensis*, which was reported to degrade 75% of low-crystallinity PET at only 28°C [7].

The identification of T*f*H, HiC, LCC, and *Is*PETase as PET degrading enzymes represent seminal works in the field of enzymatic plastic degradation, and since their discovery classical protein engineering approaches have been applied to increase their thermostability and activity [15]. T*f*H activity and thermostability has been improved from 0.043 mg_PET_/h/mg_enzyme_ at 55°C to >3 mg_PET_/h/mg_enzyme_ at 75°C [18, 15]. LCC activity and thermostability has been improved to 120 mg_PET_/h/mg_enzyme_ at 72°C in the LCC^ICCG^ variant through rational computational design [19, 18]. *Is*PETase has undergone several iterations of protein engineering to improve thermostability and thereby appararent activity. The wildtype *T*_*M*_ of 45.1°C has been increased through rational design or structural comparison with other PET hydrolases [20, 21, 22, 23, 24, 25, 26], machine learning [27], and directed evolution [24, 28, 29, 30]. The highest *T*_*M*_ was achieved in HotPETase at 82.5°C [28].

A computational method has been used to mine novel PETases by searching for sequence and structural homology between known PETases and proteins in metagenomes. This resulted in 13 potential PET hydrolase homologs, of which four were experimentally validated for PET degrading activity. [9] The method suggested by Danso et al. [9] was used to mine thermotolerant PET hydrolases from metagenomes generated based on biological material from thermal springs to ensure optimal growth temperatures >50°C [10]. In total, 37 enzymes were purified, expressed, and shown to have PET hydrolyzing activity. [10]

Here, a new approach for designing a polycarbonate (PC) degrading enzyme is presented. To this end, an active site was designed to maximize transition state stabilization of the hydrolysis of polycarbonate based on the catalytic triad of PETases. The optimized active site structure with a PC-substrate was docked into a library of highly thermostable protein scaffolds. The use of these scaffolds should ensure that the designed enzymes are stable at high temperatures and in the presence of organic solvents. The best design was produced recombinantly in *E. coli* and validated experimentally.

## MATERIALS AND METHODS

### Materials

Solvents, buffer components, and chemicals were obtained from Sigma-Aldrich, VWR International, Fluka, or Merck unless otherwise specified. All chemicals were used as supplied without further purification. Endonucleases and *E. coli* BL21(DE3) (New England Biolabs). TEV protease (Addgene, pRK793). Makrolon 2808 (gift from Covestro).

### Enzyme Design

#### PETase Homology and Catalytic Motif Selection

Multiple sequence alignment (MSA) and phylogenetic tree analysis in Clustal Omega [31] were used to compare subclasses of ester hydrolases (EC 3.1.1 and EC 3.1.2) including PETases. All proteins used had a solved structure with a resolution of <2 Å and a single chain without mutations. PET degrading enzymes were identified in the phylogenetic tree. PETases and a subset of enzymes that were evolutionarily similar were used to identify conserved catalytic motifs. Five PET degrading enzymes were identified, and a total of nine enzymes were selected. A Ser-His-Asp/Glu catalytic triad was selected as catalytic residues for the hydrolysis of polycarbonate.

Structural homology of enzymes resulting from the MSA were assessed by using the template modeling score (TM-score) defined by Zhang et al. [32], which was calculated for each cross-wise pair of proteins identified in the MSA. Structural data for 24 PETases with solved crystal structures identified in the PAZy database [6] was obtained from the protein data bank (PDB). The TM-score matrix was then used to cluster proteins using a cut-off of 0.5 with t-distributed stochastic neighbor (t-SNE) embedding [33] in Scikit-learn [34].

#### QM Validation of Catalytic Triad

Quantum mechanics (QM) simulations were used to confirm that the Ser-His-Asp/Glu catalytic triad stabilized the transition state for the hydrolysis of polycarbonate. In QM simulations, diphenyl carbonate was used as an analogue of polycarbonate. The serine, histidine, and aspartic acid in the catalytic triad were substituted by methanol, imidazole, and a formate anion, respectively. QM simulations were run in ORCA version 4.2.1 [35]. Initial and final states of the system were energy minimized with a composite PBEh-3c approach using the def2-SVP basis set [36], gCP [37], and D3BJ [38] corrections. The reaction pathway was calculated by a nudged elastic band approach [39], and the transition state along the intrinsic reaction coordinate was obtained once optimization of the reaction pathway had converged.

#### Scaffold Selection

A database of 1,302 scaffolds was generated from the PDB database based on the following criteria: scaffolds had resolutions < 2.5 Å, their M_W_ was > 13 kDa, one unique protein chain, no mutations, and either naturally thermostable or *de novo* designed.

#### Grafting the Catalytic Geometry

The catalytic triad including the oxyanion hole was grafted to scaffolds using Rosetta [40]. For a match to be generated, two criteria needed to be fulfilled: (1) it needed to have enough space for the ligand and (2) the geometry of the catalytic residues with respect to the ligand are as defined. The catalytic geometry was defined in a cst file. It was obtained by measuring the geometric constraints between the substrate, 1-(2-hydroxyethyl) 4-methyl terephthalate (HEMT), and the catalytic triad and oxyanion hole as obtained from the crystal structure of a PETases (PDB ID: 5XH3). One distance, two angles, and three dihedral restraints are necessary to completely describe the catalytic geometry between the ligand and a residue [40]. The tolerances defined in the cst file were adapted from Guanlin et al. [41]. A params file of the ligand (the PC dimer shown in Figure 2b) was generated. The catalytic geometries specified in the cst file and the ligand were matched to the scaffold database.

Rosetta was used to optimize the proteins generated. The cutoff distances used were < 8 Å for the designable region and < 12 Å for the repackable region. Three cycles of design and subsequent minimization occurred for each redesign. For each match, 25 redesigns were created. A final repack without the ligand was completed to optimize the packing of the introduced residues.

#### Design Selection

The designs were evaluated and subsequently reduced in five steps (Figure 3). (1) Designs better than the average value across all selected design parameters were chosen (Figure 4). The weighted average was defined as

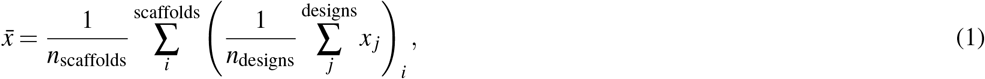

where 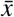 is the weighted mean, *n*_scaffolds_ is the number of scaffolds, *n*_designs_ is the number of designs for a scaffold *i*, and *x*_*j*_ is the parameter in question for a design *j*. The average used to determine the cut-off was weighted according to the number of designs per scaffold, such that the weighted average did not over-represent scaffolds with many designs. (2) Representative designs were chosen so that every design for every scaffold had unique catalytic placements or motifs. (3) Assessment and exclusion based on whether the catalytic motif coincides with the interface between a homo polymeric biological assembly. (4) Assessment of ligand binding affinity with Autodock Vina [42, 43] to obtain an empirical value of binding affinity. BPA-based polycarbonate with two and four repeating units were docked. (5) Molecular dynamics simulation of final six selected designs to determine protein-substrate interactions and distance.Molecular modeling was carried out with GROMACS [44, 45]. A Verlet cutoff scheme [46], LINCS restraint algorithm [47], particle mesh Ewald electrostatic treatment [48], and *xyz* periodic boundary conditions were used. A simulation was run for 10 ns with a step size of 1 fs using a Parrinello-Rahman barostat [49].

### Design Validation

#### Enzyme Production and Purification

The designed PC hydrolase, based on the scaffold of Y_III_(Dq)_4_CqI (PDB ID: 5MFB) [50], was produced recombinantly in *E. coli*. The protein is referred to as mutant designed armadillo repeat protein (mdArmRP). A variant of the protein without the active site (harboring mutation S108A), referred to as the wild type designed armadillo repeat protein (wtdArmRP), was used as a negative control.

The amino acid sequences were back translated using EMBOSS Backtranseq [31] with respect to codon preferences for *E. coli* K12. To facilitate easy purification, a polyhistidine tag was introduced at the 5’ end directly downstream of the translational start codon. A TEV protease cleavage site [51] was placed directly downstream of the polyhistidine tag to ensure removal of the polyhistidine tag after purification. Both genes were chemically synthesized and integrated into pET11a [52] vectors via their flanking *Nde*I and *Bam*HI restriction sites by BIOCAT GmbH. The resulting plasmids are referred to as p*mdArmRP* and p*wtdArmRP*.

Both genes were expressed in *E. coli* BL21(DE3) cells transformed with plasmid p*mdArmRP* and plasmid p*wtdArmRP*. The cells were grown in LB medium containing 100 mg/L ampicillin at 37°C and 270 rpm to an OD_600_ of 0.7, and induced with 0.1 mM IPTG for 5 hours. Cells were harvested by centrifugation at 5000 g for 20 min at 4°C, and resuspended in lysis buffer (20 mM sodium phosphate, 300 mM sodium chloride, 10 mM imidazole at pH 8), frozen, thawed, and sonicated (Sonics vibra-cell, microtip, 6 min (4 sec on/6 sec off), and amplitude of 29%). The lysed cells were centrifuged at 12,000 g for 30 min at 4°C and the supernatant was filtered (0.45 µm) and purified by affinity chromatography (HisTrap FF 5 mL, Cytiva). The column was equilibrated with buffer A (20 mM phosphate buffer, 300 mM NaCl, 10 mM imidazole at pH 7.4). A linear gradient from 0–100% buffer B (20 mM phosphate buffer, 300 mM NaCl, 250 mM imidazole at pH 7.4) was applied over 12 CV (column volumes). Elution of wtdArmRP and mdArmRP protein from the column was achieved at 20–35% buffer B. The purified protein was dialyzed against activation buffer (50 mM Tris-HCl, 0.5 mM EDTA, 1 mM DTT, pH 7.5). Polyhistidine tags were cleaved with TEV protease [51] produced recombinantly in house.

To cleave the polyhistidine tag of the wtdArmRP and mdArmRP, the TEV protease was mixed in a molar ratio of 1:100 (TEV protease to enzyme) and incubated for 3 hours at 30°C. The solution was dialyzed against buffer A, and the protein was purified with affinity chromatography (HisTrap FF 5 mL, Cytiva). The column was equilibrated with buffer A, the sample was loaded onto the column, and it was subsequently washed with 3 CV of buffer A. The flow through was collected, and the purified protein was dialyzed against ultra-pure water overnight and lyophilized.

#### Thermostability

Circular dichroism (CD) spectra were measured at room temperature on a Jasco J-715 spectropolarimeter in a 0.1 mm cell. Thermal denaturation curves were obtained by thermal scanning the sample from 10°C to 90°C at a scan rate of 2°C/min. The characteristic *α*-helical CD signal at 222 nm (*θ*_222_) was monitored, and the midpoint denaturation temperature (*T*_*M*_) was estimated by fitting a two-state model [53].

#### Electrostatic Mapping of the Surface

The electrostatic surface potential was calculated using TITRA [54] and Delphi [55]. A bulk protein dielectric constant of 4 was assumed. For water, the dielectric constant was set to 78.5, and the ion-exclusion layer was set to 2.5 Å, and the burial parameter was set to 0.4 Å. Atomic accessibility was calculated using a probe radius of 1.4 Å. The electrostatic surface potential was mapped using Delphi and visualized in VMD [56].

#### Activity Towards Polycarbonate

Polycarbonate thin films were fabricated using a spin-coater (POLOS spin-coater). A silicon wafer (1##x00D7;1 cm) was spin-coated with 3.2% (W/V) Makrolon 2808 (Covestro) dissolved in cyclopentanone at 1500 rpm and 1200 rpm/s acceleration for 90 seconds followed by annealing at 170°C for 5 minutes. The wafers were incubated at 40°C with 50 *µ*M enzyme in 50 mM Tris HCl (pH 8.0) at a final volume of 150 *µ*L. Samples were prepared with both mdArmRP and wtdArmRP, and a sample was incubated with 50 mM Tris HCl (pH 8.0) as reference. After incubation, the wafers were washed with 150 *µ*L water. The samples were analyzed using AFM (NT-MDT Ntegra) in tapping mode with NSG10 (Tips Nano) cantilevers.

## RESULTS

### Natural PETase Homology

Multiple sequence alignment (MSA) of ester hydrolases including PETases show that PETases have conserved sequence homology (Figure 1a). This is not the case when comparing PETases with other ester hydrolases. Beyond sequence homology, topology is also largely conserved for PETases (assessed for PETases for which structural data is available). This is illustrated in Figure 1b with two PETases which have a template modeling score (TM-score) of 0.65. A TM-score of > 0.5 indicates the same topology [32]. A structural evaluation of PETases also illustrates that all conform to an *α*/*β*-hydrolase fold as observed elsewhere [57]. Using t-SNE embedding with TM-scores for ester hydrolases shows a distinct cluster for most natural PETases (Figure 1c).

**Figure 1.**
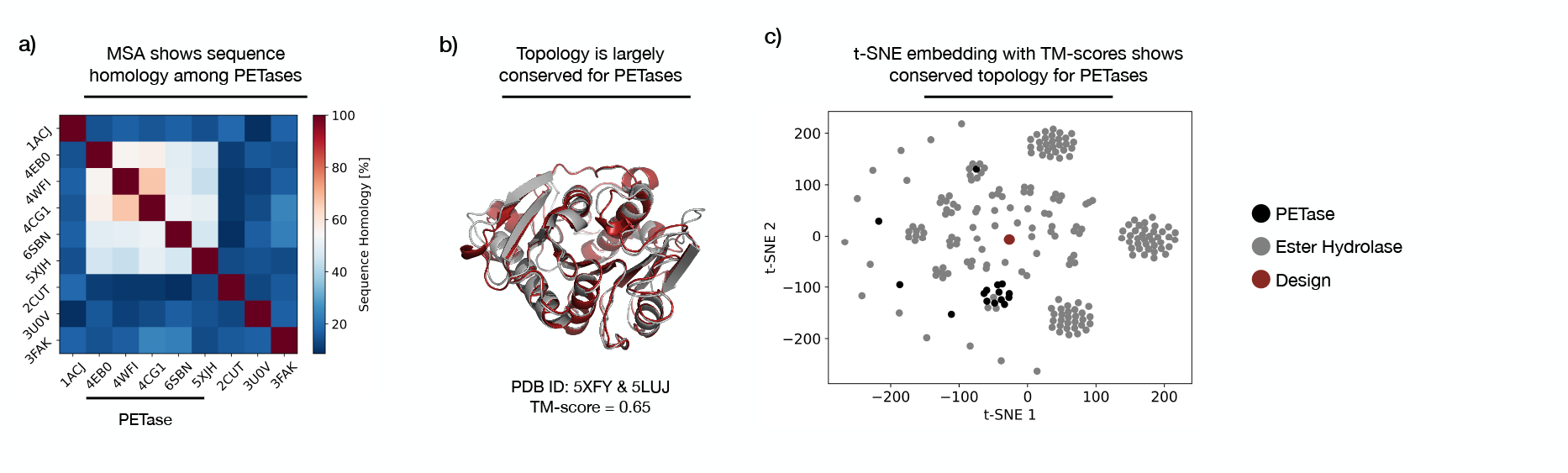
A structural investigation of PETases. a) MSA of nine selected ester hydrolases including known PETases shows increased sequence homology among PETases. The bar labeled ‘PETase’ indicates the known PETases. The enzymes are denoted by their PDB ID. b) TM-score for two PETases shows that topology is conserved. c) t-SNE embedding by TM-scores of ester hydrolases including PETases shows a distinct cluster, indicating conserved topology for most PETases.

The results indicate that PETases are strongly bound in sequence and topology, and as a consequence may be restricted in terms of achievable stability and catalytic efficiency, potentially limiting optimization. Indeed, directed evolution and rational design of PETases to increase melting temperature show diminishing returns in terms of activity [24, 28, 30]. It is thus of interest to design enzymes with activity towards plastics from highly thermostable scaffolds with a low homology to natural PETases as to escape the sequence space by which they appear to be confined.

### Active Site of PETases

Conserved for all the enzymes depicted in Figure 1a is a Ser-His-Asp/Glu triad. The angles and distances between the catalytic residues, for the seven enzymes with a Ser-His-Asp triad, along with the distances between the residues for the Ser-His-Glu triad, show that the geometry is preserved. Figure 2a depicts this phenomenon for the five identified PETases. From Figure 2a it is evident that even down to the backbone trace the active site is conserved. This implies that this conserved motif is important for catalysis. Thus, to use the knowledge nature has provided, a Ser-His-Asp/Glu triad is used as an outset to design a catalytic geometry for PC degradation.

**Figure 2.**
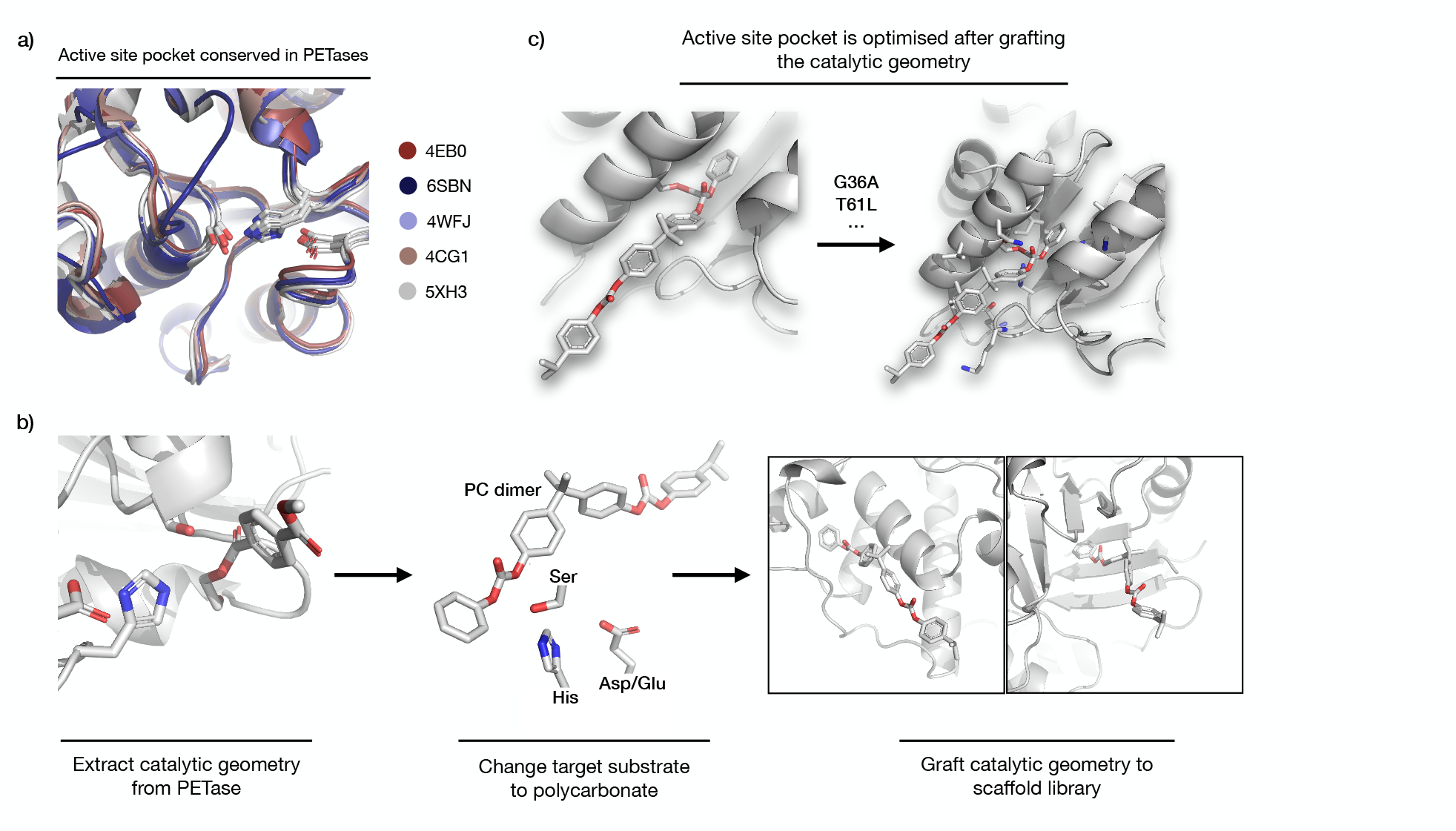
Methodology for grafting the catalytic geometry into a scaffold library. a) Shows the active site pocket is conserved across PETases. The enzymes are denoted by their PDB ID. b) Illustrates the catalytic geometry from PETase (PDB ID: 5XH3) to which HEMT is bound. The catalytic geometry is extracted and the ligand is exchanged with the target substrate polycarbonate. The new catalytic geometry is grafted into a scaffold library. A polycarbonate (PC) dimer is chosen as the desired substrate. c) After grafting the catalytic geometry, the binding pocket is optimized often resulting in mutations to hydrophobic residues.

### Defining the Catalytic Geometry for Grafting

For one of the identified PETases, a solved crystal structure of the protein with a ligand bound was identified (PDB ID: 5XH3). The catalytic geometry for PC degradation was obtained from the distances, angles, and dihedral restraints of the PETase with a bound ligand (HEMT). Quantum mechanical simulations of the transition state of a Ser-His-Asp triad with a substrate analogue (diphenyl carbonate) resembles that of the bound ligand with respect to the catalytic triad in the PETase. While the ligand bound state is not a transition state, the quantum mechanical simulation encourages its use as such (data not shown). Furthermore, the catalytic triad is shown to stabilize the transition state for polycarbonate, thereby facilitating rate acceleration of the reaction (data not shown). It therefore seems reasonable to use the catalytic geometry obtained from a ligand bound crystal structure for the design of a polycarbonate degrading enzyme. The methodology is illustrated in Figure 2b.

Grafting the catalytic geometry, including the target substrate, ensures an appropriate pocket is identified which facilitates the catalytic geometry. The scaffold library ensures thermostability, which is essential for enzymatic activity near the glass transition temperature of plastics. The active site pocket may be further optimized by mutation to confer higher binding affinity by introducing complementary interactions with the substrate, as well as stabilizing the desired geometry (Figure 2c).

While a hydrolase catalytic triad may not be the ideal solution for polycarbonate hydrolysis, this was an attempt to verify whether PC degrading activity could be implanted in an inactive scaffold. Theoretically, numerous catalytic arrangements can be screened *in silico*, using QM simulations to identify transition state stabilizing geometries [58]. A catalytic triad, however, has a well understood mechanism and has been documented to have promiscuous hydrolase activity [59]. Both PET and PC have a carbonyl group in common, which is the target of the nucleophilic attack.

#### Design Exclusion

The design methodology using Rosetta resulted in 27,034 designs across 160 scaffolds, and the process used for selection and evaluation of the designs is visualized in Figure 3. Step (1) evaluates the catalytic interaction and scaffold integrity using selected design parameters from the Rosetta score file. The selected design parameters are mentioned in Figure 4, which also illustrates how many designs pass the cut-off for each parameter and the total number of designs that pass the cut-offs for all selected design parameters. Step (1) of evaluation resulted in 169 designs across 19 scaffolds.

**Figure 3.**
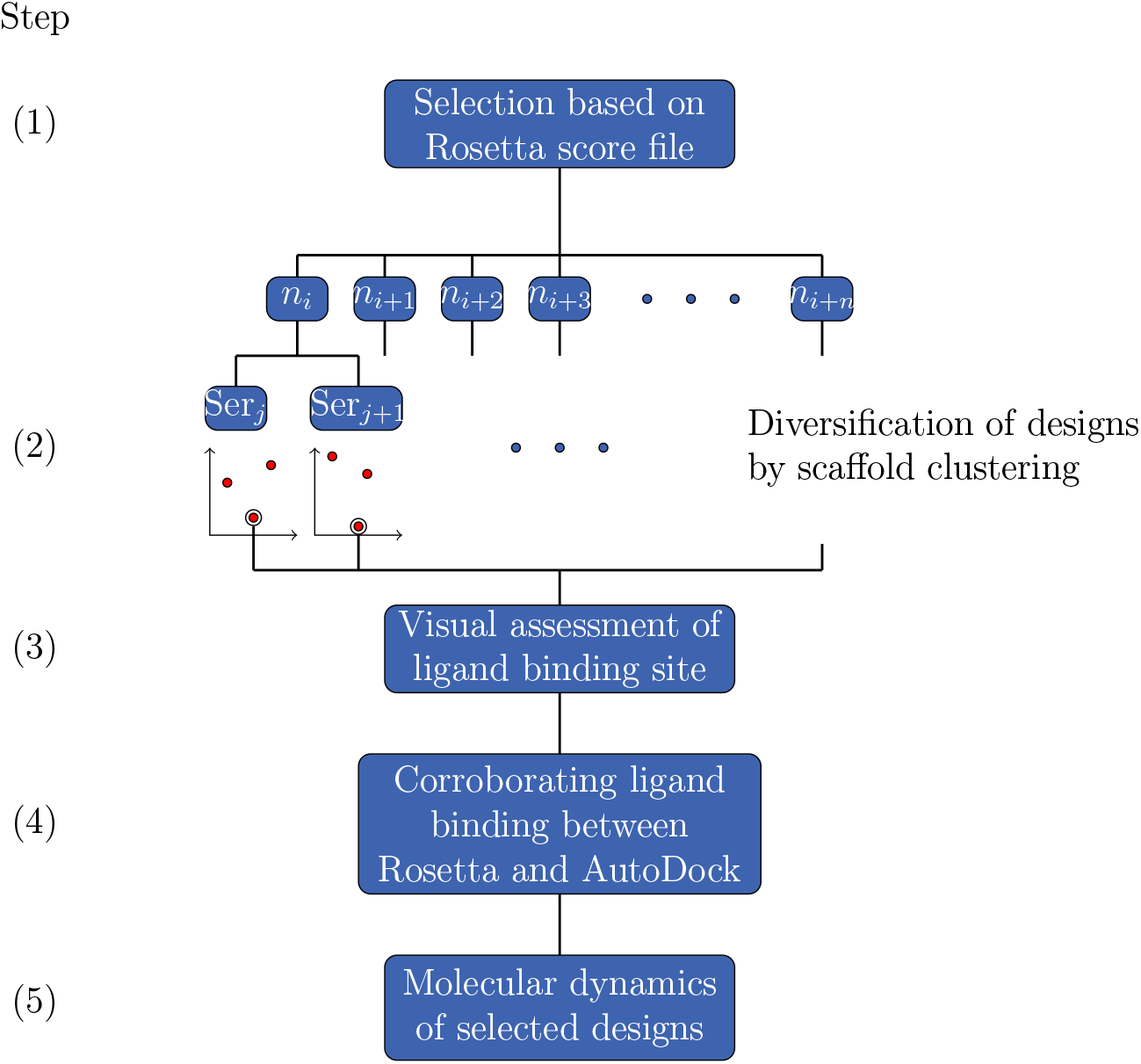
The flowchart visualizes the five step evaluation procedure used to find optimal designs. Step (1) consists of calculating the weighted mean (Equation 1) for selected design parameters and picking designs, with a score in the top 50th percentile for all selected design parameters. These were then ordered according to ligand binding score. Step (2) consists of a clustering by scaffold and then subsequent catalytic residue clustering. For each catalytic residue cluster one representative design was picked. Step (3) consists of rational evaluation by assessing whether the binding interface coincides with the protein interface in polymeric proteins. Step (4) was to assess ligand binding affinity with Autodock Vina to obtain an empirical value of binding affinity. Step (5) consists of molecular dynamics simulation of final selected designs to determine protein-substrate interactions and distance.

**Figure 4.**
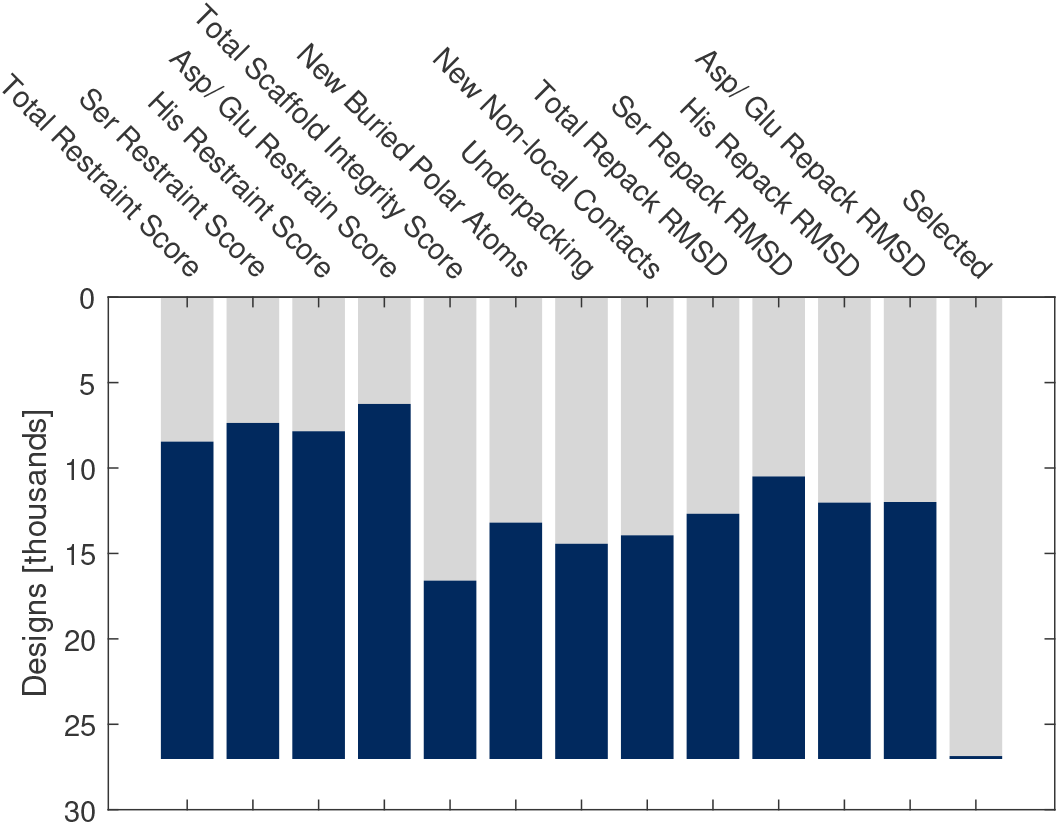
The plot shows the number of designs that passed the cutoff for the selected design parameters used to evaluate the designs. The last bar to the right named ‘Selected’ shows the total number of designs that passed all selected design parameters. The blue bar represents the designs that pass the cutoff and the gray bar represents the designs that did not pass the cutoff. ‘Ser Restraint Score’, ‘His Restraint Score’, ‘Asp/Glu Restraint Score’, ‘Ser Repack RMSD’, ‘His Repack RMSD’, ‘Asp/Glu Repack RMSD’, and ‘Total Repack RMSD’ are measures for the catalytic geometry. ‘Total Restraint Score’, ‘Total Scaffold Integrity Score’, ‘New Buried Polar Atoms’, ‘Underpacking’, and ‘New Non-local Contacts’ are measures for the scaffold integrity.

The parameters from the Rosetta score file only consider the catalytic geometry, which is necessary for transition state stabilization, and the scaffold integrity, which is necessary for protein stability. However, ligand-protein interactions are also necessary to ensure the formation of the enzyme-substrate complex [60].

To compensate for the missing weighting of protein-ligand interaction in step (1) of evaluation, ligand affinity was used for evaluation in step (2). In step (2), the designs are clustered according to different catalytic residues in a single scaffold, and then the design with best ligand affinity and scaffold integrity score is chosen. This resulted in 33 designs across 19 scaffolds. Choosing the best design for each scaffold with either a unique catalytic placements or motif ensures a more diverse subset of designs while still reducing the number of designs. The choice of the design with the highest ligand affinity might make product release problematic, however, ligand affinity is important to assure that an enzyme-substrate complex will form [60].

Scaffold integrity ensures that the designed protein is not less stable than the thermostable scaffold. Highly stable proteins are desirable for plastic degradation as high temperatures increase reaction rate and make the polymer chains more accessible. [14, 13, 60]

In step (3), the position of the active site is evaluated. Designs with the active site placed in an interface of a homo polymeric biological assembly are excluded. This type of active site placement hinders access of the substrate to the active site. Step (3) yielded 19 designs across 13 scaffolds.

#### Computational Evaluation of Potential PC hydrolases

In step (4), AutoDock Vina was used to check whether the binding site and mode matched with those predicted by Rosetta. If they did not match, the designs were sorted out, which reduced it to six designs. Generally, AutoDock Vina and Rosetta results agree in predicting the binding sites for the final six designs across five scaffolds as seen in Figure 5. There is, however, variation due to the allowed torsional angles (rotable bonds between quaternary carbon and aromatic rings) for polycarbonate in AutoDock Vina, which in some cases allow the substrate to wrap around the surface (PDB ID: 1CLC, 5CSR, 5MFB). Furthermore, the highest affinity conformer does not always coincide with the Rosetta result. In all cases, the distances between polycarbonate and serine are increased when polycarbonate is docked with Autodock Vina. The angles between polycarbonate and serine are, however, similar for both docking results.

**Figure 5.**
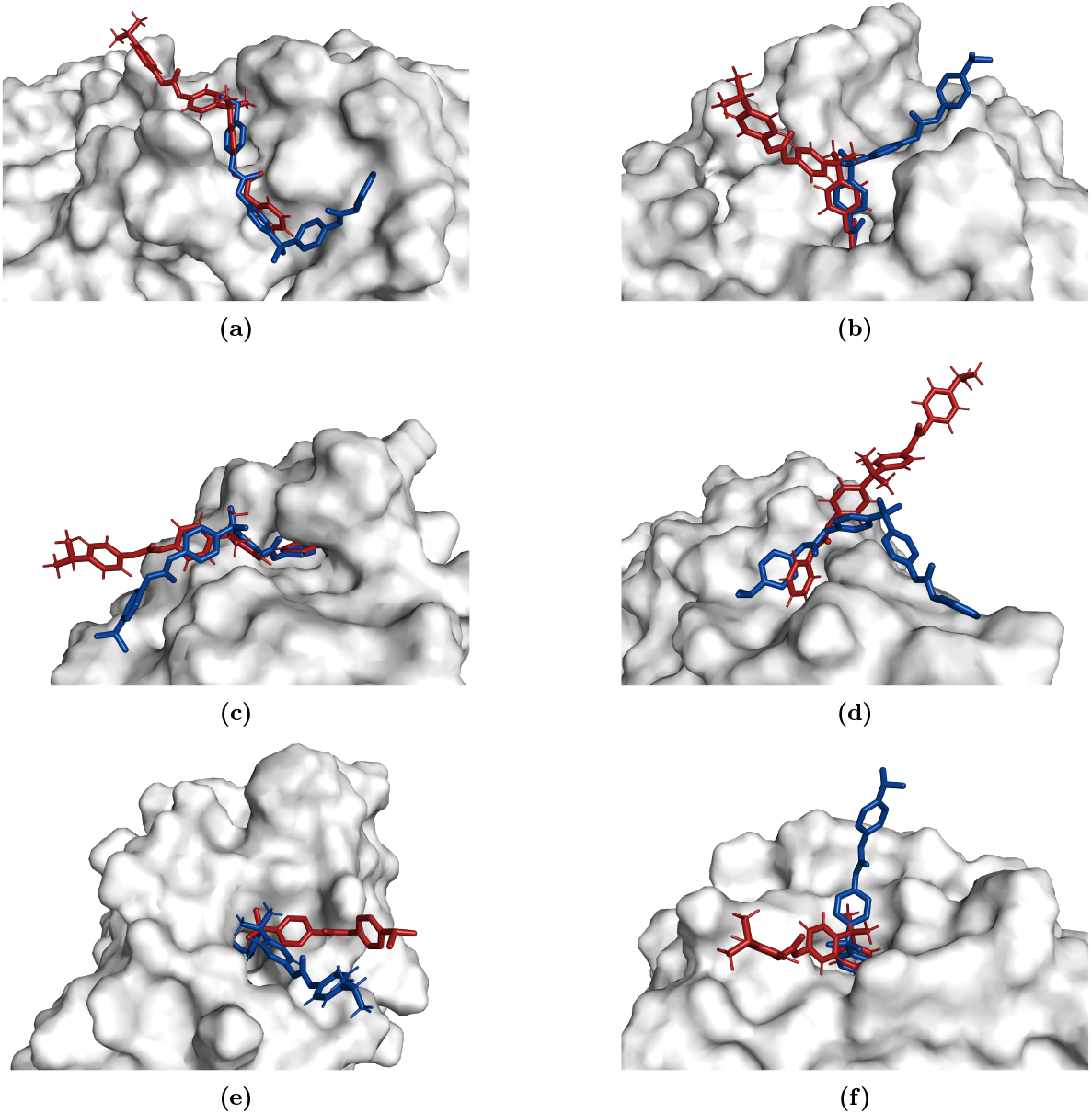
The active sites of the six potential PC hydrolases visualized with ligands docked using Rosetta in red and ligands docked using AutoDock Vina in blue. The PDB IDs of the scaffolds of the visualized designs are (a) 1CLC, (b) 5CSR, (c) 5FQK, (d) 5MFB, (e) 6C7H* with catalytic residues S65, H51, E53, and N68, and (f) 6C7H** with catalytic residues S65, H51, D53, and N68. The surface of the enzyme is colored gray.

The three-dimensional shape of the binding pockets influences the binding affinity as the designs with deeper active sites have higher binding affinities compared to the binding affinities of designs with a shallow active site cleft. The embedding of a substrate in a deep pocket also allows for more protein-substrate interactions, yielding specific binding. This suggests that a deeper pocket contributes both higher binding affinity and more specific binding. However, polycarbonate in bulk form is an aggregated substrate, and as such, the enzyme designed must be able to catalyze the hydrolysis of aggregated substrates. It has been shown that PET degrading enzymes often have an open, shallow active site [61]. For an aggregated substrate, the polymer chains are packed densely, allowing for little fluctuation of the chains. As a result, it might be necessary for the active site to be open and shallow to reach the polymer chains as is the case for some designs (PDB ID: 5MFB and 1CLC).

To further examine the substrate specificity, a polycarbonate tetramer was also docked into the chosen designs using AutoDock Vina. For the 5MFB design (mdArmRP), the tetramer wraps around the protein surface, and the carbonate group docks in the same place as seen for the polycarbonate dimer (Figure 6c). This was not the case for designs with deeper active site pockets, which indicates that the shallow active site cleft of mdArmRP is favorable.

**Figure 6.**
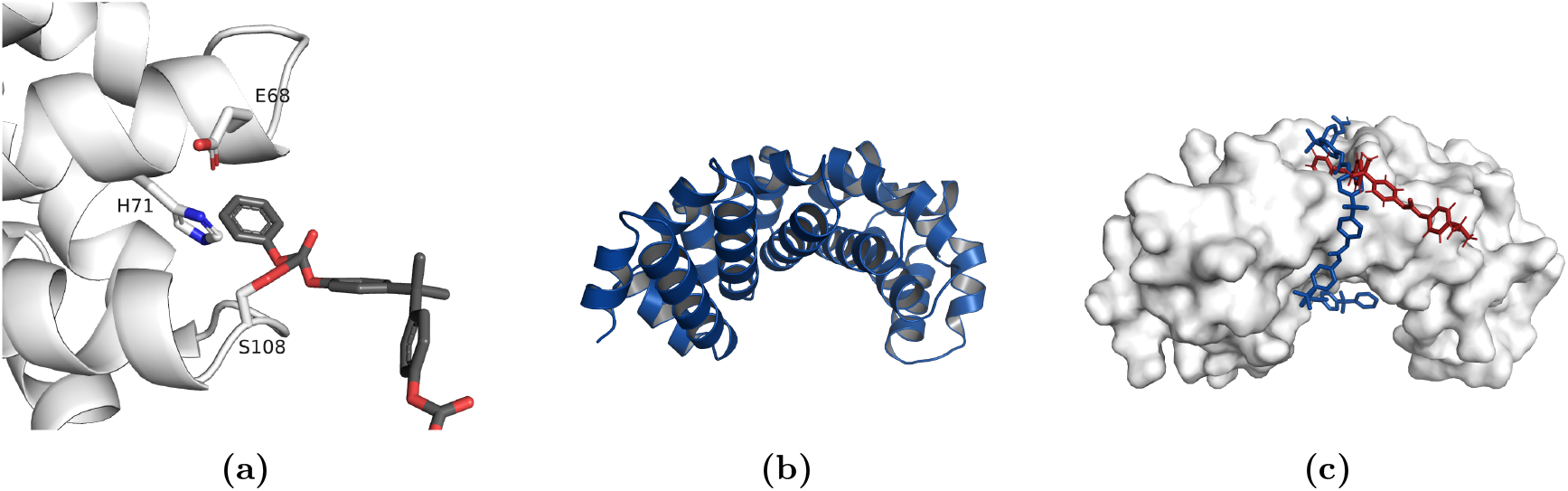
(a) Visualization of the active site with a docked polycarbonate dimer of the designed PC hydrolase, mdArmRP. The catalytic residues shown in blue are S108, H71, and E68. (b) Predicted structure of the mdArmRP. The structure is predicted by Rosetta. (c) Visualization of the designed PC hydrolase, mdArmRP, with a polycarbonate dimer docked using Rosetta in red, and a polycarbonate tetramer docked using AutoDock Vina in blue. The tetramer is a repeating structure of bisphenol A. The ligand affinity for the docked polycarbonate tetramer is -10.3 kcal/mol.

Molecular dynamics simulations were used to evaluate catalytic occupancy and hydrogen bonding in step (5). The Ser-PC distance was used to evaluate catalytic occupancy, and while the sampling of catalytically relevant distances (2.86 Å as defined in the cst file) does occur for the selected designs, the catalytic occupancy is minimal with respect to the 10 ns molecular dynamics simulation. This indicates that enzymatic activity is likely low, as the frequency of sampling conformations with relevant distances is low. Static protonation states were used for MD simulations, and hence, serine was protonated in the simulation. As a result, the corresponding distances expected from molecular dynamics between serine and the substrate should be comparably larger. Trajectory data also shows that polycarbonate remains within 8 Å of the serine through the entire simulation.

The MD simulation showed that for many of the designs the mean hydrogen bonding between polycarbonate and the designed proteins is less than one. This implies that non-polar contacts are the basis of ligand binding, which is coherent with the limited number of polar contacts possible with polycarbonate.

Overall, the MD simulation showed that all six designs performed similarly, and therefore, the choice of a final PC hydrolase design was based on the Autodock Vina results and the number of mutations to the scaffold. A low number of mutations is desirable to minimize their potential destabilizing effect. As a result, the design based on the 5MFB scaffold (mdArmRP) was chosen for experimental validation.

#### Evaluation of Final PC Hydrolase

The armadillo repeat protein (PDB ID: 5MFB) is a synthetic construct [50]. The fold of mdArmRP is shown in Figure 6b. In order to implement the active site into this scaffold only four mutations to the original scaffold protein were necessary. Notably, the catalytic serine is already present in the scaffold, thereby increasing the chances of activity of the final design further. A histidine is introduced and has the effect of increasing the reactivity of the catalytic serine. The general trend of the introduced mutations is a loss of charge and the introduction of smaller residues. A loop region and an *α*-helix are used to position the catalytic triad, this is shown in Figure 6a.

The electrostatic surface potential of mdArmRP was analyzed at pH 4, 7, and 9 to evaluate at which pH enzymatic activity is expected. The catalytic histidine needs to be deprotonated for the catalytic triad to catalyze the reaction [62]. The electrostatic surface potential is visualized in Figure 7. This shows that at pH 4, the active site has a positive electrostatic surface potential which indicates that the catalytic histidine is protonated. The surface potential of the catalytic histidine shifts from positive to negative when pH is increased from 7 to 9. A titration curve was plotted for the catalytic histidine, which shows that the histidine is deprotonated above pH 8 (data not shown). This indicates that a pH above 8 would be favorable for catalysis.

**Figure 7.**
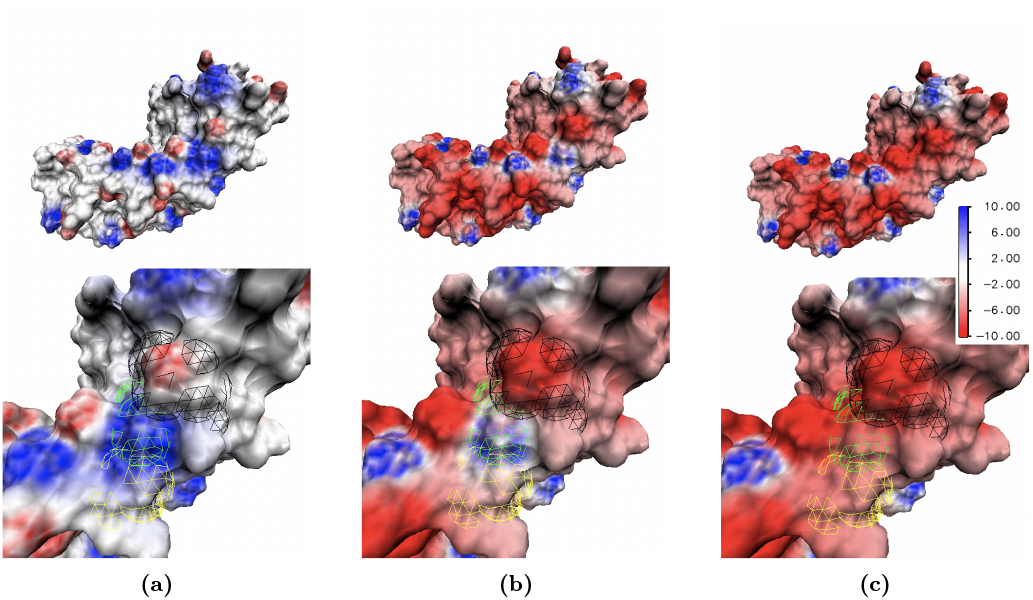
The electrostatic surface potential in units of *k*_*B*_T/e of mdArmRP at (a) pH 4, (b) pH 7, and (c) pH 9. The upper images show the full protein while the lower images are a zoom of the active site where the catalytic residues are marked with a colored mesh. Serine is marked with yellow, histidine is marked with green, and glutamic acid is marked with black. The titration profile was calculated with TITRA [54], mapped with Delphi [55], and visualized with VMD [56].

An alternating pattern of positive and negative charges observed on the surface of mdArmRP is the result of a Glu/Lys salt bridge motif. The salt bridges may contribute to the high thermostability of the protein. The salt bridges are retained within the investigated pH range from pH 4 to pH 9. The high proportion of helical secondary structure further ensures many intramolecular hydrogen bonds, which contribute to the overall high thermostability. Interactions between helices are both hydrophobically and electrostatically driven by charge-charge interactions of the salt bridges. The net contribution of these parameters results in a highly stable protein.

#### Experimental Validation of PC Hydrolase

Thermostability of the protein is analyzed using CD spectroscopy, and the *T*_*M*_ for mdArmRP is estimated to be 102°C. This was, however, predicted as described in the method, which means that it is an estimate and not the exact *T*_*M*_. The *T*_*M*_ of the wild type is not reported, but the *T*_*M*_ of similar armadillo repeat proteins is around 50–90°C [63]. This indicates that the introduced mutations do not destabilize the protein. A high *T*_*M*_ is favorable for catalysis and accessibility to aggregated substrates. CD spectroscopy also showed that the (*θ*_222_) is almost fully restored after partial unfolding, which means that mdArmRP refolds completely after being heated to 90°C. Reversibility of unfolding is important as this assures that the enzyme can be reused for multiple rounds of plastic degradation.

Polycarbonate hydrolysis activity was tested and confirmed using AFM. As seen in Figure 8 there is significant variation on the surface of the polycarbonate film incubated with buffer compared to the PC hydrolase, which implies polycarbonate degradation. Crystalline regions, which have not been degraded, are seen in the surface incubated with PC hydrolase. This suggests that the PC hydrolase can degrade amorphous polycarbonate but is unable to access crystalline patches. Activity was not detectable for the wtdArmRP (data not shown).

**Figure 8.**
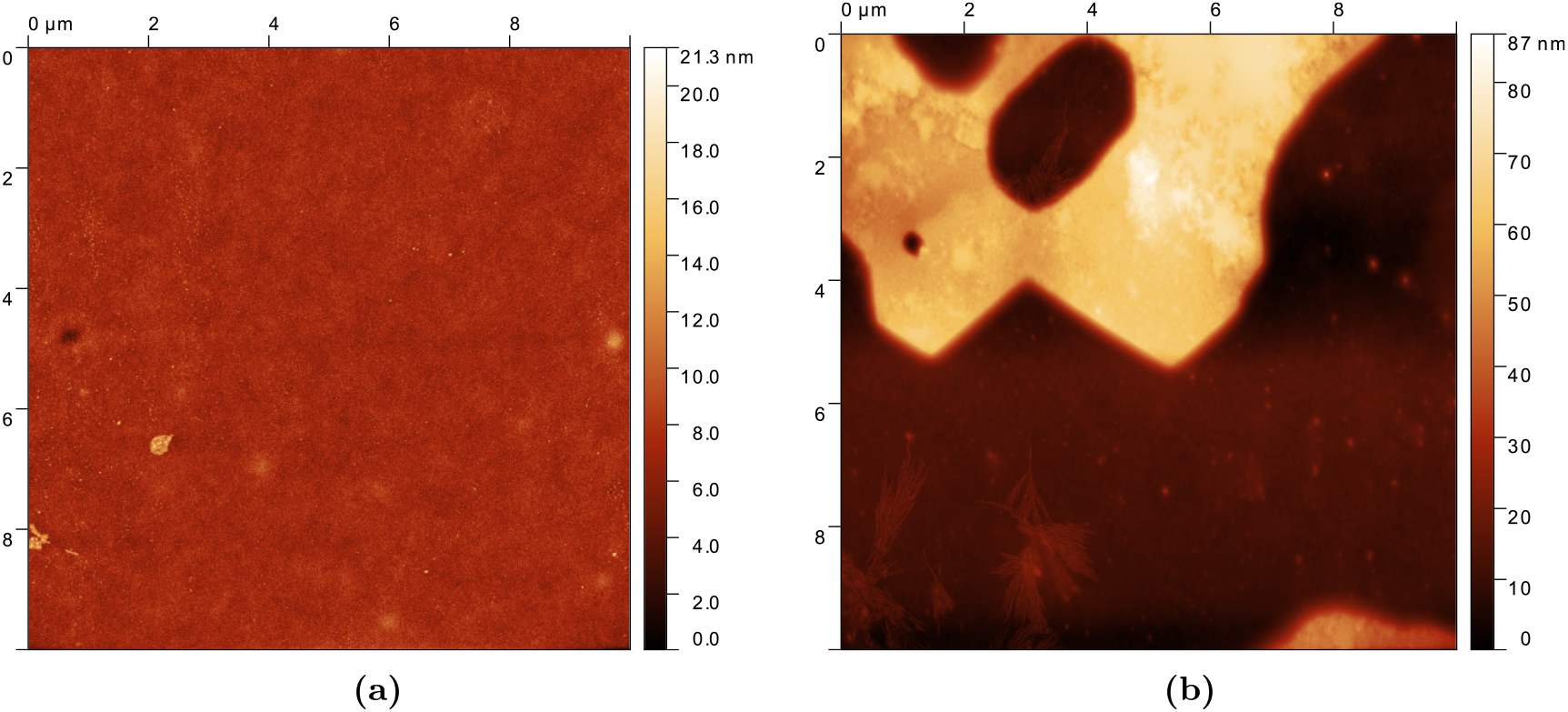
10 × 10 *µ*m AFM images of polycarbonate films spin coated onto silicon substrate. The polycarbonate films were incubated at 40°C for 96 hours with (a) 50 mM TrisHCl at pH 8, and (b) with 40 mM mdArmRP in 50 mM TrisHCl at pH 8.

## CONCLUSIONS

The results from modeling and experimental validation serve as a proof-of-concept on the utility of enzyme redesign as a method to introduce catalytic activity into thermostable scaffolds to produce a plastic degrading enzyme. This has the advantage of escaping the sequence space, which is occupied by natural proteins and offers a new starting point for directed mutagenesis. This method can be used to engineer stable enzymes with activity towards plastics.

